# Genetically engineered MRI-trackable extracellular vesicles as SARS-CoV-2 mimetics for mapping ACE2 binding *in vivo*

**DOI:** 10.1101/2022.03.27.485958

**Authors:** Andrea Galisova, Jiri Zahradnik, Hyla Allouche-Arnon, Mattia I. Morandi, Paula Abou Karam, Ori Avinoam, Neta Regev-Rudzki, Gideon Schreiber, Amnon Bar-Shir

**Affiliations:** Department of Molecular Chemistry and Materials Science, Weizmann Institute of Science, Rehovot, 7610001, Israel; Department of Biomolecular Sciences, Weizmann Institute of Science, Rehovot, 7610001, Israel

## Abstract

The elucidation of viral-receptor interactions and an understanding of virus-spreading mechanisms are of great importance, particularly in the era of pandemic. Indeed, advances in computational chemistry, synthetic biology, and protein engineering have allowed precise prediction and characterization of such interactions. Nevertheless, the hazards of the infectiousness of viruses, their rapid mutagenesis, and the need to study viral-receptor interactions in a complex *in vivo* setup, call for further developments. Here, we show the development of biocompatible genetically engineered extracellular vesicles (EVs) that display the receptor binding domain (RBD) of SARS-CoV-2 on their surface as coronavirus mimetics (EVs^RBD^). Loading EVs^RBD^ with iron oxide nanoparticles makes them MRI-visible, and thus, allows mapping of the binding of RBD to ACE2 receptors non-invasively in live subjects. Importantly, the proposed mimetics can be easily modified to display the RBD of SARS-CoV-2mutants, namely Delta and Omicron, allowing rapid screening of newly raised variants of the virus. The proposed platform thus shows relevance and cruciality in the examination of quickly evolving pathogenic viruses in an adjustable, fast, and safe manner.

## Introduction

Virus-receptor recognition is the initial step in the infectious cycle and is considered to be a key stage in the induction of viral pathogenesis^1^. Therefore, the elucidation of the interactions of viruses with host cells’ receptors is of paramount importance for a better understanding of pathology pathways and for the development of antiviral interventions. For example, it has been shown that the severe acute respiratory syndrome coronavirus 2 (SARS-CoV-2), which has caused the current and prolonged coronavirus disease 2019 (COVID-19) pandemic, specifically attacks cells expressing high levels of angiotensin-converting enzyme 2 (ACE2) ^2^. The understanding of the interactions between the receptor binding domain (RBD) of the spike-S protein of the virus with ACE2 ^3^ has resulted in the development of a wide range of efficient therapeutics and vaccines ^4–8^. Unfortunately, SARS-CoV-2 has shown a remarkable ability to rapidly introduce mutations to the spike protein and the RBD for improved affinity and immune evasion ^9–12^, which have led to rapid spreading of more transmissible variants and compromised effectiveness of available vaccines ^13^. Thus, it is clear that there is a need for the ability to characterize viruses and their evolving mutants quickly and safely and potentially even predict dangerous variants before they emerge. Indeed, *in silico* ^14^ and *in vitro* ^15^ examination of viruses provides crucial insights into virus-receptor interactions. Nevertheless, these approaches are limited in the study of off-target binding events and are not applicable for spatial and real-time mapping of viral-receptor binding in deep tissues. This calls for a method with the ability to longitudinally and non-invasively monitor and map *in vivo* viral distribution and receptor binding in a safe and rapid way to enhance our ability to study emerging viruses and assess biological feedback to therapeutics.

Several types of non-viral nano-sized formulations have been proposed to elucidate viral-receptor interactions so far, including those for studying SARS-CoV-2 ^4,16–18^. Among these, extracellular vesicles (EVs) offer several advantages over synthetic nanoparticles. First, as cellular content nanocarriers ^19–21^, they are biological substances, suggesting that they can be introduced into the body without leading to the side effects often encountered with synthetic formulations. Second, they can be genetically engineered to present biomolecules on their surface, providing a rapid and general method for the display of peptides ^22–24^. As such, EV targetability to a tissue of interest has been enhanced by displaying peptides that are not present on the surface of native EVs ^25–28^. Third, EVs share important similarities with enveloped viruses, including comparable sizes and host membrane compositions ^29,30^. EVs are thus attractive non-infectious vehicles with which to exploit viral uptake pathways for cellular cargo delivery ^25,31–33^, or for the development of EV-based vaccines ^34^ and related adjuvants ^35^. For example, EVs presenting the coronavirus S protein or its RBD were proposed as potential vaccines already in the mid-2000s for SARS-CoV ^36^, as well as for the current SARS-CoV-2 pandemic ^37,38^. Moreover they showed efficiency as decoys for neutralizing antibodies ^39^ and as systems for targeted delivery of antiviral agents ^40^. In addition, the ease at which EVs can be genetically engineered makes these formulations ideal for rapid studies of emerging viral mutations as they appear.

Given that EVs can mimic viruses and can be labeled with imageable material ^27,41,42^, EVs can potentially be used for non-invasive *in vivo* imaging of viral-receptor interactions. In fact, it has been shown that EVs can be fluorescently labeled and imaged *in vivo;* however, these fluorescent methods are unable to track EVs in deep tissues and offer limited spatial resolution ^43^. In contrast, tracking of EVs with threedimensional imaging modalities (such as CT ^41^ and MRI ^42^) allows an assessment of their spatial distribution even in deep tissues. In this regard, MRI stands out due to its ability to provide spatial information from the introduced EVs that can be overlaid on high-resolution anatomical images of the same subject avoiding the need of using hybrid multimodal imaging approaches. Here, we show the design, development, and implementation of genetically-engineered EVs that display the RBD of SARS-CoV-2 (EVs^RBD^) as coronavirus mimetics for studying RBD-ACE2 interactions. Magnetically-labeled EVs^RBD^ allow mapping RBD-ACE2 binding *in vivo* and in real-time using a clinically translatable MRI setup. Moreover, we demonstrate the modifiability of the EV-based formulation by presenting the RBD of currently spreading SARS-CoV-2 variants - Delta and Omicron. The ability to monitor both *in vivo* biodistribution and the effect of different binding affinities of RBD to ACE2 in a fast and safe way highlights the potential of our approach in prolonged pandemic eras and for the study of other emerging viruses.

## Results and Discussion

### Genetic design of SARS-CoV-2 receptor binding domain (RBD) constructs

Several methods have already been implemented to genetically engineer EVs to display peptides on their surface as, e.g., in fusion with the Lamp2b EVs membrane protein ^32,44^, the vesicular stomatitis virus G protein (VSVG) ^40^, or the transmembrane domain of platelet-derived growth factor receptor (PDGFR) when using the pDisplay™ vector ^27^. Although widely used, Lamp2b showed an inability to express the RBD of the SARS-CoV-2 on the membrane efficiently ^40^. Starting from the pDisplay™ vector, which has been extensively used for heterologous expression and surface display of cell receptors in mammalian cells, we first engineered it for efficient expression of viral peptides on the surface of EVs. To this end, and with a purpose to create SARS-CoV-2 mimetics - EV^RBD^ (Figure 1), we constructed a new pAGDisplay plasmid through a three-component assembly of fragments from the widely used pDisplay, pLVX-Puro, and pET26b plasmids. There are four main benefits of using our designed pAGDisplay plasmid over other alternatives. First, it uses the puromycin resistance marker, which is a more potent selective antibiotic compared to geneticin. Second, an IRES sequence, which was introduced downstream from the transmembrane domain to allow co-expression of the antibiotic resistance and the transmembrane domain under a single promoter restricting expression quenching. Third, the fluorescent protein eUnaG2 was fused with the C-terminus of PuroR, allowing for easy detection of transfected cells. Fourth, we introduced a RFnano protein, a MIRFPnano670 derivative (see the Materials and Methods section), at the C-terminus of the PDGFβ transmembrane domain to allow efficient selection of RBD-expressing cells using fluorescent activated cell sorting (FACS). Having designed the pAGDisplay plasmid for efficient and versatile surface display of cell receptors, the Wuhan RBD of SARS-CoV-2 was first cloned at the N-terminus preceding the transmembrane PDGFR domain. Specific tags were added to the obtained constructs for further validation of expression with an ALFA tag as a marker to the RBD construct and a Myc-tag to the control construct (referred to later in the text as noRBD).

**Figure 1.**
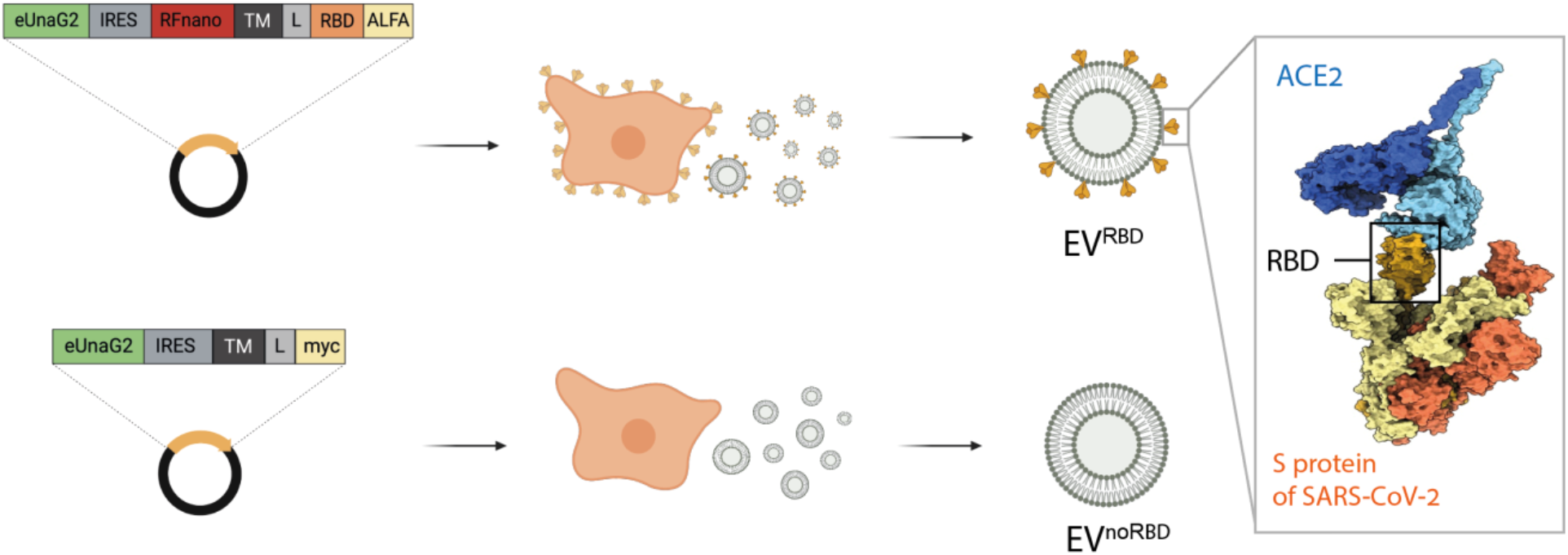
Schematic illustration of the proposed design. From left to right: A scheme of the pAGDisplay plasmid designed for this study. The released EVs display RBD (EV^RBD^) and control EVs display no RBD (EV^noRBD^) from HEK293 cells transfected with pAGDisplay-RBD and pAGDisplay-noRBD, respectively. On the right, a model of interaction between the RBD of the spike protein of the SARS-CoV-2 virus and the ACE2 receptor; RBD is highlighted in the black square). eUnaG2 – a green fluorescent protein; IRES – internal ribosomal entry site; RFnano – red fluorescent protein (MIRFPnano670); TM – transmembrane domain; L – linker; RBD – receptor binding site of SARS-CoV-2; ALFA – ALFA tag; Myc – Myc tag.

### Parental cells display functional RBD with wild type affinity to ACE2

Human embryonic kidney 293 (HEK293) cells were transfected with pAGDisplay-RBD or pAGDisplay-noRBD followed by FACS to select cells associated with the highest expression levels of eUnaG2 (green fluorescence) and RFnano (red fluorescence). Stable cell lines expressing RBD or controls were established under puromycin antibiotics selection for three weeks. HEK293 cells stably expressing RBD on the membrane surface (RBD cells) or control cells (noRBD cells) were obtained and characterized using immunostaining, immunoblotting, flow cytometry (Figure 2a-c) and confocal microscopy (Figure S1). The presence of RBD fused to the ALFA tag on the cell surface was confirmed using a designed anti-ALFA tag nanobody (DnbALFA, see the Materials and Methods section) fused with mNeon Green protein (excitation 506 nm, emission 517 nm). RBD-expressing cells incubated with the nanobody showed a bright signal on the cell surface corresponding to the presence of RBD, in contrast to noRBD cells, which showed only a background signal of the eUnaG2 protein present in the cytoplasm (Figure 2a). The binding of the anti-ALFA tag nanobody only to the RBD cells was also confirmed at a single-cell level by flow cytometry (Figure 2b, p < 0.001 and Figure S2 for FACS data). Moreover, Western blot analysis of lysed cells showed the expression of the RBD-ALFA tag protein only in RBD cells (Figure 2c).

**Figure 2.**
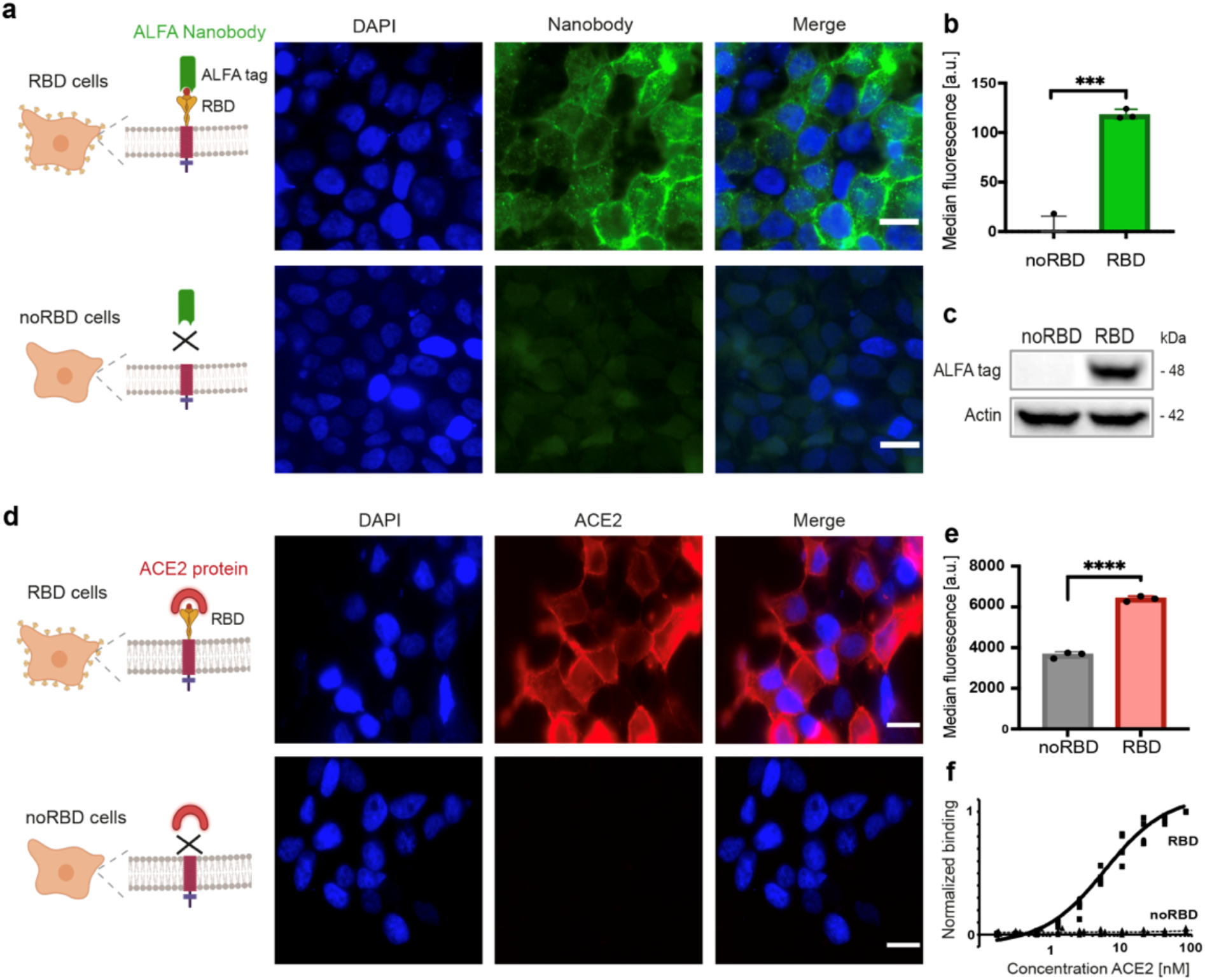
Genetically engineered cells expressing RBD on their surface. **(a)** Schematics of the experimental design (left) and representative fluorescence images of HEK293 cells expressing RBD (RBD cells, top) or control cells (noRBD cells, bottom) after incubation with anti-ALFA tag nanobody (green). DAPI (blue) stained cell nuclei. **(b)** Flow cytometry quantification of fluorescent signals from RBD and noRBD cells labeled with the anti-ALFA tag nanobody (n=3, p-value=0.0002). **(c)** Western blot analysis of lysates of RBD and noRBD cells. ALFA tag fused with RBD protein resulted in a 48 kDa protein; actin served as a housekeeping protein. **(d)** Targeting of cells (RBD cells, top; noRBD cells, bottom) with fluorescently conjugated ACE2 protein (schematically shown at the left), and fluorescent images of cells incubated with fluorescently labeled ACE2 protein (red). Blue represents the cells’ nuclei (stained with DAPI). Red fluorescence of RFnano is below the set threshold. The scale bar represents 20 μm. **(e)** Median fluorescence (from flow cytometry experiments) of RBD cells and noRBD cells incubated with the ACE2 protein (n=3, p-value=0.00001). **(f)** Binding curves of ACE2 to RBD and noRBD cells quantified by flow cytometry. Data are presented as mean values ± s.d. Statistics: two-tailed unpaired Student’s t-test with ***p-value<0.001, and ****p-value<0.0001.

Having demonstrated the presence of RBD on the cell surface, we then tested its functionality. For this purpose, the recombinant extracellular portion of ACE2 protein was expressed, purified, and conjugated to a fluorophore CF640R (excitation 642 nm, emission 662 nm) followed by its incubation with either noRBD or RBD cells (Figure 2d-f). Fluorescence microscopy images showed a clear binding of the ACE2 protein only to RBD cells as a higher fluorescent signal compared to controls, confirming the presence of a functional RBD on the cell surface (Figure 2d). Similarly, flow cytometry analysis showed a significantly higher fluorescent signal in RBD cells compared to noRBD cells (p < 0.001), corresponding to the bound ACE2 protein (Figure 2e and Figure S2 for FACS data). By incubating the cells with a range of different concentrations of fluorescently labeled ACE2 protein and performing a series of FACS experiments to determine its affinity to RBD at the cell surface, an equilibrium dissociation constant (*K*_D_) of 6.7 ± 1.3 nM was determined for RBD cells (Figure 2f). This affinity of RBD expressed at the surface of HEK293 cells to the ACE2 protein is in accordance with the values reported for purified proteins or through a yeast-display assay ^10,45^.

### EVs^RBD^ as functional SARS-CoV-2 mimetics that target ACE2 receptors

Following the validation of the expression of functional RBD at the surface of HEK293 cells through ACE2 binding assessments (Figure 2), secreted EVs of these cells were obtained using a standard method of differential centrifugation of the cells’ media (with an average 5×10^9^ EVs/ one million cells) according to MISEV guidelines ^46^. The isolated and purified EVs from RBD cells or control cells were referred to as EVs^RBD^, or EVs^noRBD^, respectively, and were further examined. Western blot analysis confirmed the expression of RBD (Figure 3a) via detection of ALFA-tag only in EVs^RBD^, while CD81 and Alix, typical markers of EVs ^46^, were detected in both EVs^RBD^ and EVs^noRBD^ (Figure 3b). Cryo-Transmission electron microscopy (TEM) images revealed the spherical morphology and typical size of vesicles for both EVs^noRBD^ and EVs^RBD^ (Figure 3c). Nanoparticle tracking analysis showed a homogenous distribution of size, with an average size of 80.9 ± 1.8 nm and 90.4 ± 1.6 nm for EVs^RBD^ and EVs^noRBD^, respectively (Figure 3d), comparable to the size reported for SARS-CoV-2 ^47^.

**Figure 3.**
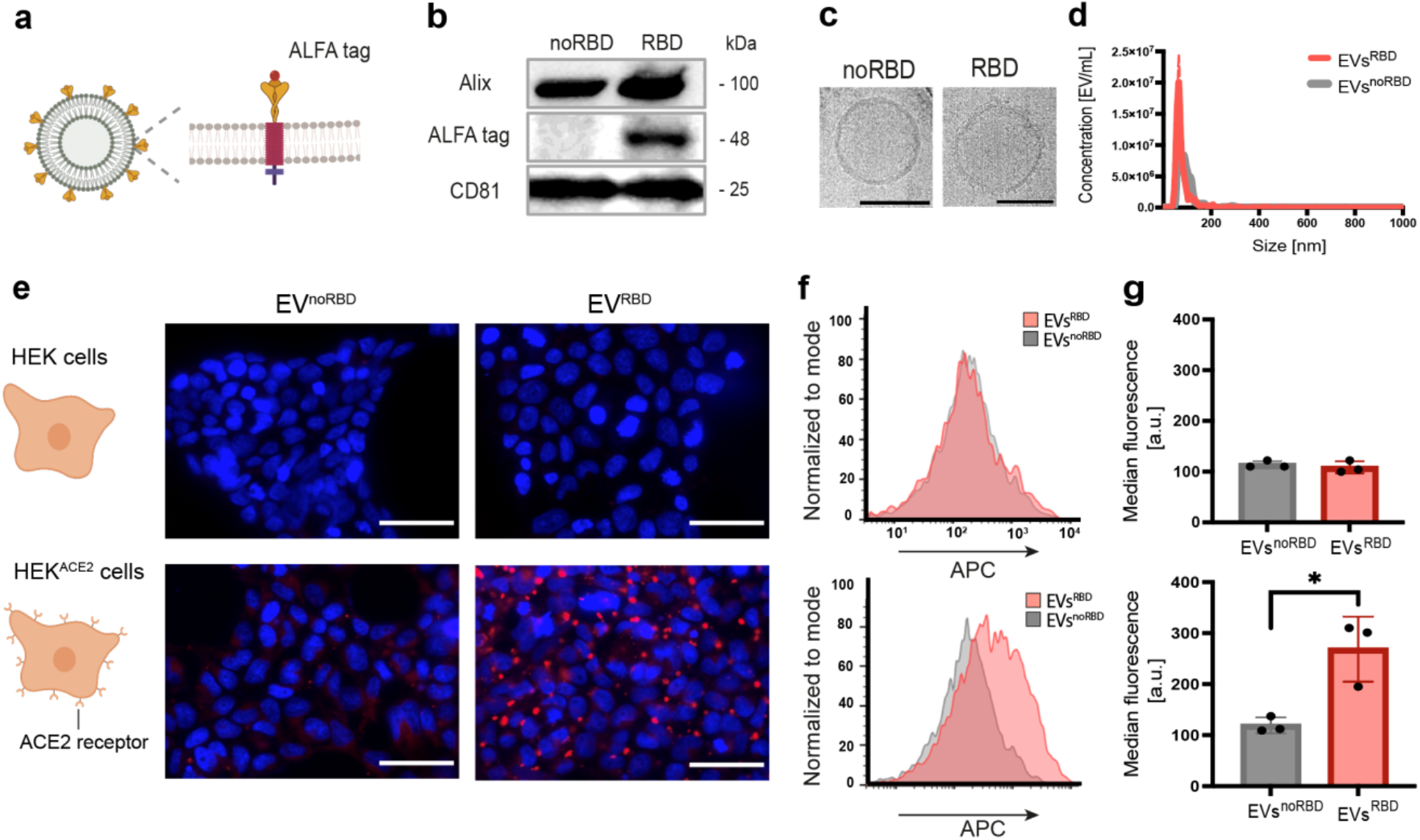
EVs as SARS-CoV-2 mimetics. **(a)** A schematic illustration of the presence of the ALFA tag on the EVs surface. **(b)** Western blot analysis of ALFA tag and CD81 expressed in EVs^noRBD^ and EVs^RBD^. **(c)** Cryo-TEM image of representative EV^noRBD^ and EV^RBD^. Scale bars represent 100 nm. **(d)** Nanoparticle tracking analysis of EVs^noRBD^ and EVs^RBD^ depicting the size of the purified EVs. **(e)** Fluorescent images of HEK293 cells (HEK, top) or HEK293 cells stably expressing ACE2 receptors (HEK^ACE2^, bottom) incubated with red-fluorescently labeled EV^noRBD^or EV^RBD^ for three hours; DAPI was used for cell nuclei staining and is shown in blue. The scale bar represents 50 μm. Flow cytometry analysis of cells following their incubation with fluorescently labeled EVs^noRBD^ and EV^RBD^ shown as histograms **(f)** and median of fluorescence intensity (n=3, p-value=0.0171) **(g).** Data are presented as mean values ± s.d. Statistics: two-tailed unpaired Student’s t-test with *p-value<0.05.

Next, with the aim to examine the binding capabilities of EVs^RBD^ to ACE2-expressing (HEK^ACE2^) as compared to control (HEK) cells, both cell types were incubated with fluorescently labeled EVs for three hours followed by their examination with fluorescence microscopy (Figure 3e) and flow cytometry (Figure 3f-g). The expression of the ACE2 receptors in the inoculated HEK^ACE2^ cells was confirmed by immunoblotting, FACS analysis and enzymatically (Figure S5). Fluorescent images of the examined control cells (HEK) showed no pronounce red fluorescence following their incubation with either EVs^RBD^ or EVs^noRBD^ (Figure 3e, top row). In contrast, a pronounced red fluorescence was obtained for HEK^ACE2^ after their incubation with fluorescently labeled EVs^RBD^, confirming their extensive binding to ACE2-expressing cells (Figure 3e, bottom row). Note here that the very low red fluorescence found in control cells after their incubation with the EVs^RBD^ may be in accordance to the expected low expression levels of native ACE2 in HEK293 cells. Flow cytometry analysis showed that EVs^RBD^ had significantly greater binding efficiency to ACE2-expressing cells (HEK^ACE2^) when compared to EV^noRBD^ (p < 0.05), while the uptake into control cells (HEK) was similar for both EVs^RBD^ and EV^noRBD^ (Figure 3f-g and Figure S3 for FACS data). These results thus confirm the presence and functionality of the engineered RBD peptide on the EVs^RBD^ surface and its targeting and binding capability to ACE2 receptors expressed on cells with no observable effect on their viability (Figure S4). These findings show that the engineered EVs^RBD^ can be used as SARS-CoV-2 mimetics with ACE2 binding capabilities with a similar size and shape to the SARS-CoV-2 virus. Importantly, this EV-based formulation, in contrast to a highly infectious SARS-CoV-2 virus ^48^, can be used to study RBD-ACE2 interactions at a single-cell level under standard laboratory conditions without the need for strict safety regulations. This platform may, therefore, expand the research of viral-receptor interactions to research institutes and industrial setups that do not yet have access to dedicated facilities, which must be designated with the highest biosafety level required to work with SARS-CoV-2.

### Biodistribution and targeting of SARS-CoV-2 mimetics in mice

The *in vivo* targetability of the EVs^RBD^ mimetics toward ACE2 receptors was then studied in an animal model. Several studies have already shown the ability of engineered EVs to interact and bind to receptors of interest *in vivo*, including acetylcholine receptors in the brain ^32^, tumor cells ^27^, immune cell surfaces^49^, and viral receptors, including ACE2 ^40^. Since human ACE2 receptors are not naturally present in mice, a few transgenic mice models had been proposed ^50,51^, but they are not readily accessible, and therefore, are not yet extensively used. Therefore, we established a xenograft model to study the *in vivo* targetability of EVs^RBD^ to ACE2 following their systemic administration. For this purpose, control HEK293 cells (HEK) and ACE2-expressing HEK293 cells (HEK^ACE2^) were first inoculated, bilaterally and subcutaneously in two flanks of immunodeficient mice to result in tumor-like appearances at the implantation sites two weeks after cell inoculation. In contrast to the transgenic mouse model ^40^, the model we used here allows for real-time comparison of the accumulation of EVs following their administration in both the control and ACE2-expressing cells, simultaneously in the same animal. Two weeks after cell transplantation, mice were injected, intravenously, with fluorescently labeled EVs, either non-targeted EVs (EV^noRBD^, group 1, n=5) or ACE2-targeted EVs (EV^RBD^, group 2, n=5) as schematically depicted in Figure 4a. Six hours after EVs administration (3×10^11^ EVs per mouse), the two types of tumors were excised for further quantitative analysis of their fluorescent (DiR) signal. Importantly, although introduced systemically (through intravenous injection), a clear and statistically significant difference in the fluorescence of HEK and HEK^ACE2^ tumors was obtained only after the administration of EVs^RBD^ with a higher accumulation in the ACE2-expressing cells (Figure b, c; p < 0.01; n = 5/group). Importantly, no observable difference in the fluorescence of the tumors could be detected after the injection of EVs^noRBD^, indicating the high targeting ability and enhanced specificity of EVs^RBD^ to the ACE2 receptors *in vivo* (Figure 4b, c, Figure S6).

**Figure 4.**
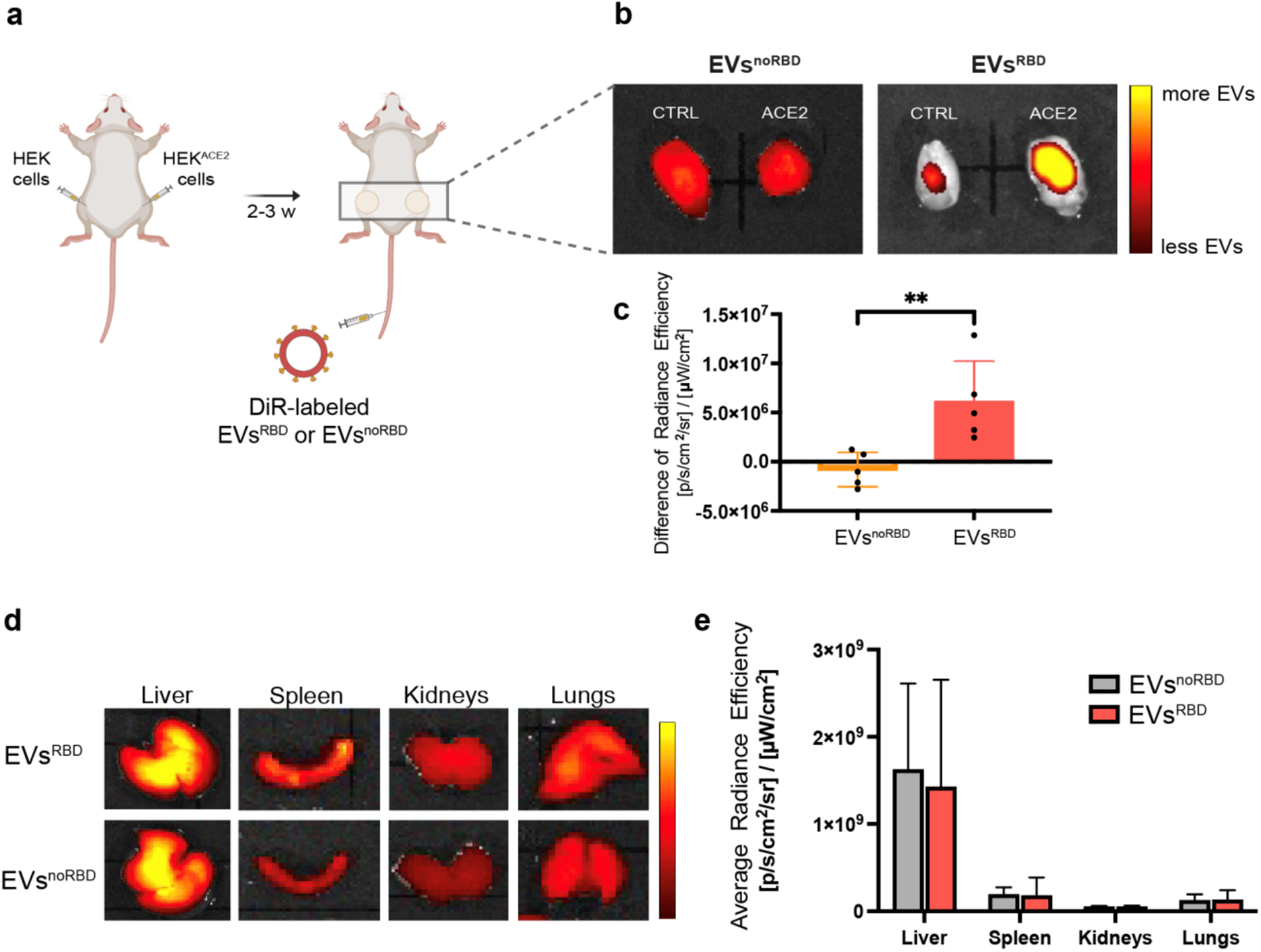
*In vivo* ACE2 targetability of EVs^RBD^. **(a)** Schematic illustration of the experimental setup at which HEK and HEK^ACE2^ cells were injected subcutaneously into the flanks of immunodeficient mice followed by intravenous administration of EVs^RBD^ or EV^noRBD^. **(b)** Representative fluorescent images of excised HEK and HEK^ACE2^tumor-like tissues six hours after intravenous administration of EVs^noRBD^ or EV^RBD^ (3×10^11^ EVs/mouse). **(c)** Quantification of the difference in the fluorescent signals obtained fromHEK and HEK^ACE2^tumors following the injection of EVs^noRBD^ or EV^RBD^ (n=5/group; p-value=0.009). **(d)** Representative fluorescence images of different organs after injection of EVs^noRBD^ and EVs^RBD^. **(e)** Quantification of the biodistribution of EVs^RBD^ and EVs^noRBD^ after their intravenous injection (n=5/group). Data are presented as mean values ± s.d. Statistics: two-tailed unpaired Student’s t-test with **p-value<0.01.

Next, we studied the biodistribution of fluorescently DiR-labeled EVs after their intravenous administration in a dose of 3×10^11^ EVs per mouse. Both types of EVs, EVs^RBD^ and EVs^noRBD^, were detected mostly in the liver (Figure 4d and 4e and Figure S7), in good agreement with previous studies demonstrating that this is the main organ through which EVs are cleared from the body^24^. In contrast to the liver, much milder fluorescent signals were also obtained from excised spleen and kidney, implying that other clearance pathways might be involved in EVs biodistribution, but with no significant difference when comparing EVs^noRBD^ and EVs^RBD^. As no human ACE2 receptors are expressed in the lungs of mice ^52^, the fluorescent signals of excised lungs following the injection of either EVs^noRBD^ or EVs^RBD^ was found to be similar and very low, as expected. Overall, the comparable biodistribution profiles of EVs^noRBD^ and EVs^RBD^ (Figure 4d and 4e) indicate, once again, the specific targetability of EVs^RBD^ to ACE2-expressing cells even *in vivo* after their systemic administration (Figure 4a and 4b).

### MRI mapping of magnetically labeled SARS-CoV-2 mimetics

After showing the specific targetability of EVs^RBD^ to ACE-expressing cells both *in vitro* (Figure 3) and *in vivo* (Figure 4), we aimed to examine the ability to map the accumulation in their target cells using a non-invasive and three-dimensional imaging modality, such as MRI, which was demonstrated to be applicable for monitoring of magnetically labeled EVs *in vivo* ^42,53–55^. To this end, and to load EVs with superparamagnetic iron oxide nanoparticles (SPIONs), parental HEK-293 cells stably expressing the RBD (Figure 1) were incubated for 24 hours with SPIONs added to their culture medium (40 μg of iron per mL). Then, the incubating medium was replaced with exosome-free medium and the released EVs^RBD^ were collected and purified by differential centrifugation (Figure 5a) and further characterized. Incorporation of SPIONs into the secreted EVs^RBD^ was clearly visualized by cryo-TEM as multiple hypointense clusters inside the lumen of the EVs (Figure 5b). Solutions containing different concentrations of SPIONs-labeled EVs^RBD^ were then examined by MRI and showed concentration-dependent T2*-weighted MRI contrast (Figure 5c). Importantly, labeling of EVs with SPIONs did not compromise their size (84.8 ± 4.3 nm vs. 94.8 ± 3.4 nm, Figure 5d), charge (−18.8 ± 3.9 mV vs. −16 ± 1.0 mV, Figure 5e), or the expression of the RBD (Figure 5f).

**Figure 5.**
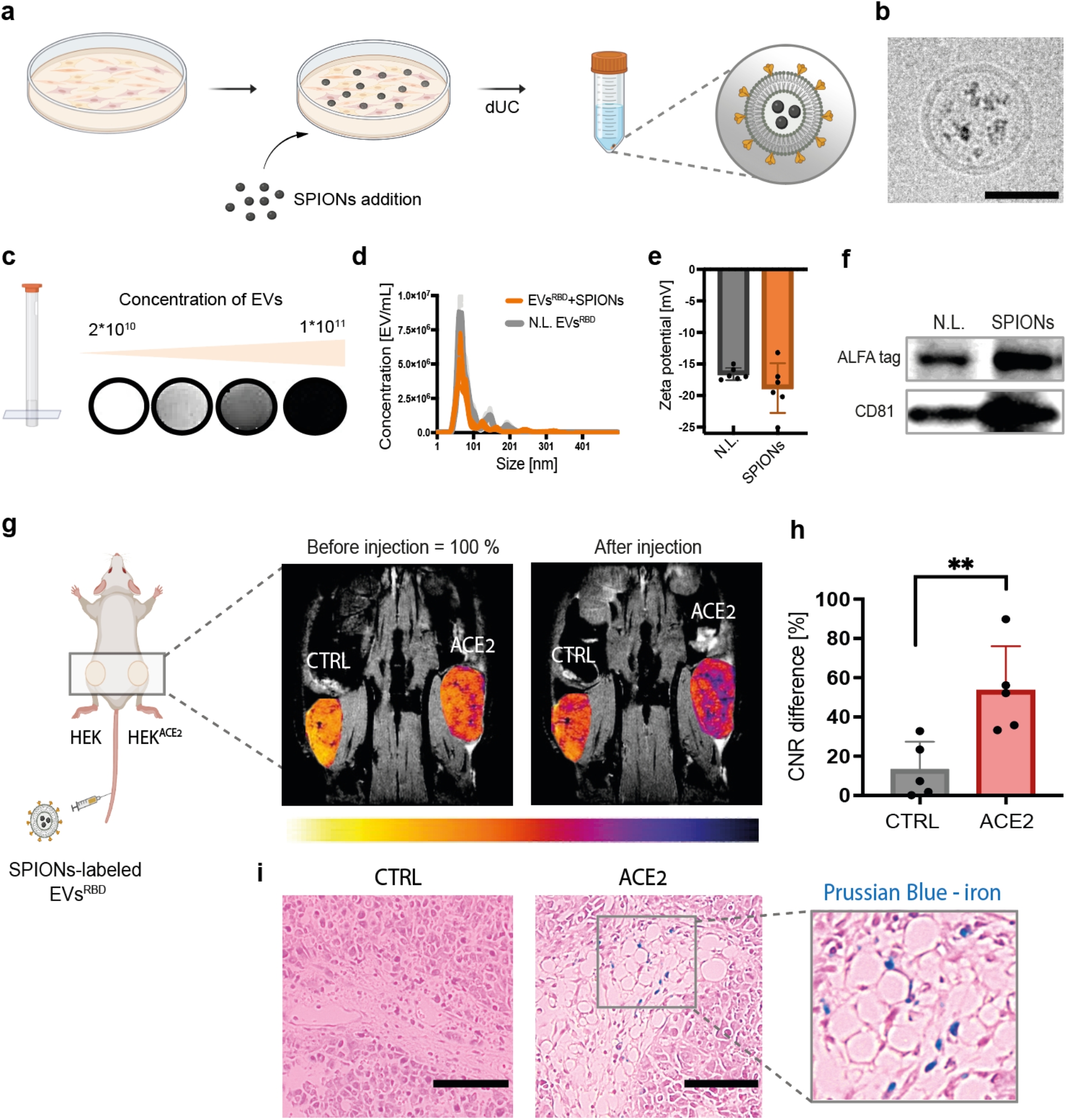
MRI mapping of ACE2 targeting of magnetically labeled EVs^RBD^. **(a)** A schematic illustration of the incorporation of SPIONs into EVs by labeling their parental cells. dUC – differential ultracentrifugation. **(b)** Cryo-TEM image of EVs with incorporated SPIONs. Scale bar represents 50 nm. **(c)** MRI of SPIONs-labeled EVs^RBD^ obtained for solutions containing different EV concentrations. **(d)** Nanoparticle tracking analysis, **(e)** Zeta potential, and **(f)** Western blot analysis of SPIONs-labeled and nonlabeled EVs. **(g)** MR images obtained before (left) and four hours after (right) the injection of EVs^RBD^ (3×10^11^ EVs/mouse). Examined tumors: control (CTRL) or HEK^ACE2^, and color-coded images of tumors overlaid on anatomical MR images of mice (gray). **(h)** Quantification of contrast-to-noise ratios (CNR) in control and ACE2 tumors after EVs^RBD^ injection (n=5, p-value=0.009). CNR was calculated as the ratio between the signal of the tumor ROI and a muscle ROI. CNR before injection was set at 100%. **(i)** Histological analysis of tumor tissues after Prussian blue staining for iron deposition (blue color). Scale bar represents 100 μm. The inset shows a magnified image of the tissue with accumulated iron in blue. Data are presented as mean values ± s.d. Statistics: two-tailed unpaired Student’s t-test with *p-value<0.05 and **p-value<0.01.

After successful labeling of the EVs with SPIONs, we aimed to examine the MRI-detectability of their ACE2 targetability *in vivo*. To this end, a solution containing 3×10^11^ SPIONs-labeled EVs^RBD^ was injected systematically through the mouse tail vein. The preferential accumulation of the SARS-CoV-2 mimetics in ACE2-expressing tissue was observed as a reduced MRI signal intensity in T2*-weighted images (Figure 5g). To quantify the change in the MRI signal of the two tumorous-like tissues (HEK *vs*. HEK^ACE2^), mice were scanned before and four hours after EVs administration and the difference in the tissue contrast-to-noise ratio (CNR) obtained before and after injection was calculated. As shown in Figure 5h, a significant increase in (p-value < 0.01, n=5) CNR difference (before vs. after SPIONs-labeled EVs^RBD^ injection) was obtained in the region of interest (ROI) of the ACE2-expressing cells compared to control tissue. Moreover, Prussian blue staining of slices of the excised tissues clearly showed iron deposits only at the samples obtained from ACE2-expressing cells (Figure 5i), confirming the delivery of the SPIONs-labeled EVs^RBD^ to HEK^ACE2^, but not to the controls.

Note here that other strategies for labeling EVs should be considered in the future to improve their iron content load, and thus, the sensitivity of the approach. Labeling secreted EVs through labeling of their parental cells might have suffered from low labeling capacity ^42,53^ and more advanced labeling methods should be developed and employed to allow the detection of lower numbers of targeted EVs in future studies. In addition to MRI mapping of the targetability of magnetically labeled SARS-CoV-2 mimetics to ACE2-expressing cells, our results confirm the successful delivery of an intravesical cargo (nanoparticles cargo) to a target tissue. The demonstration that engineered RBD-tagged EVs with encapsulated siRNA can be used to suppress SARS-CoV-2 infection in a transgenic mouse model of ACE2 expression ^40^ suggests that our platform can be used to deliver several types of cargos in the future, including antiviral or immunosuppressive agents to ACE2-expressing tissue. As such, the proposed SARS-CoV-2 mimetics could be of high importance during the current SARS-CoV-2 pandemic, where efficient and safe treatment strategies against COVID-19 are still needed.

### SARS-CoV-2 mimetics—a versatile tool to study viral binding

As for other RNA viruses, so for SARS-CoV-2, the rapid evolution and the introduction of diverse mutations at the RBD manipulates not only the binding affinity to the host cell receptors and the infectivity of the virus, but also reduces the efficiency of proposed vaccines and other therapeutics. In that regard, one key feature of the SARS-CoV-2 mimetics presented here is that suspicious mutations or those found in rapidly spread variants of a virus of interest can be easily displayed at the surface of the EVs through a one-step cloning procedure into the pAGDisplay plasmid. This means that the method can provide rapid data on novel mutations as they appear. To examine this, we have designed and tested several mimetics that represent different binding affinities to the ACE2 receptor, namely the Wuhan, Delta and Omicron variants of SARS-CoV-2 (Figure 6 and Supplementary Sequences).

**Figure 6.**
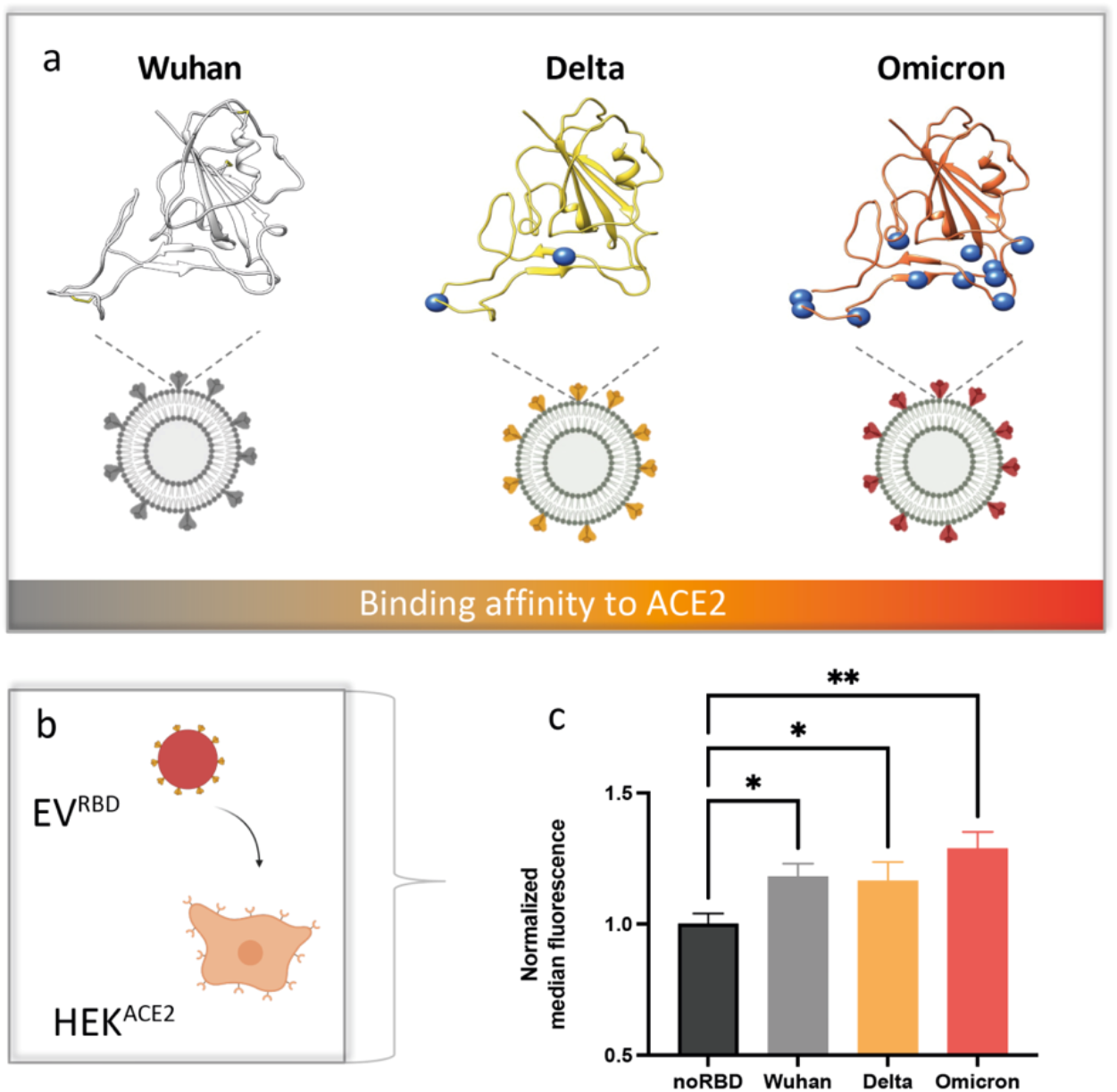
EVs representation of SARS-CoV-2 mutants. **(a)** Location of the RBD domain mutations characteristic for the Wuhan, Delta and Omicron variants of SARS-CoV-2 depicted in the Wuhan structure (pdb 6m17:f); the variations from the Wuhan variant are shown as blue circles. **(b)** Schematic illustration of the experiment depicting the binding of EVs^RBD^ to ACE2 expressing at the surface of cells. **(c)** Flow cytometry analysis of ACE2-expressing cells (n=3) incubated with fluorescently labeled EVs: EVs^noRBD^, EVs^RBD-Wuhan^, EVs^RBD-Delta^ or EVs^RBD-Omcron^. Data are presented as mean values ± s.d. Statistics: two-tailed unpaired Student’s t-test with *p-value<0.05 and **p-value<0.01.

The genes encoding for the mutated peptides were designed and cloned into the expression plasmid and the parental cells were transiently transfected. After EVs isolation to obtain EVs^RBD-Wuhan^, EVs^RBD-Delta^ and EVs^RBD-Omicron^, the difference in their binding capabilities to ACE2 receptors expressed at the surface of live cells was examined using flow cytometry. As shown in Figure 6, and as expected, HEK^ACE2^ cells incubated with engineered EVs displaying the Delta variant of RBD showed similar fluorescence when compared to cells incubated with the Wuhan RBD, implying on similar binding capabilities. In contrast, HEK^ACE2^ cells incubated with EVs presenting the Omicron variant on their surface (EVs^RBD-Omicron^) significantly higher fluorescence was obtained, as expected for RBD with a stronger binding affinity to ACE2 ^56,57^. These results demonstrate that the proposed genetically engineered EVs, which are here used as coronavirus mimetics, can be used as a reliable platform for fast and safe examination of evolved mutations of SARS-CoV-2, with the potential to be extended to other viruses.

## Conclusion

In summary, we showed the design and implementation of genetically engineered EVs as SARS-CoV-2 mimetics (EVs^RBD^), which efficiently bind the ACE2 receptor both *in vitro* and *in vivo*, even after intravenous systemic delivery. The ability to load EVs^RBD^ with MRI-trackable material (SPIONs) allowed mapping of their targetability in live subjects and thus opens new possibilities to study viral-receptor interactions in deep tissues, such as the lungs, which might have played a significant role in viral infectivity and are not accessible to luminescent-based imaging approaches. Having demonstrated the modifiability of the approach and proposing EVs that mimic additional variants of SARS-CoV-2 show its potential to be applied for currently evolving mutants in a safe and reproducible manner without the risk of using infectious material. We envision that the demonstrated strategy could be employed to study other virus-receptor interactions, beyond SARS-CoV-2-ACE2 demonstrated here, by simply engineering a tailored peptide on the EVs surfaces and using them in a wide spectrum of viral-induced pathologies. Finally, with the increasing interest in EVs research, beyond viral mimetics, including their use as cellular-derived nanocarriers, the reported visualization strategy could be of high cruciality to enrich our understanding of EV targetability to the organ of interest in the context of a live organism.

## Acknowledgments

This project received funding from the following sources: European Research Council (ERC-StG no. 677715 to A.B.-S.), the Israel Science Foundation (ISF no. 1637/20 to O.A, N.R.-R. and A.B.-S. and 1268/18 to JZ and GS), the Clore Institute for High-Field Magnetic Resonance Imaging and Spectroscopy (A.B.-S.), the Helen and Martin Kimmel Institute for Magnetic Resonance Research (A.B.-S.), and the Ben B. and Joyce E. Eisenberg Foundation (to A.B.-S. and O.A). A.B.-S. is the incumbent of the Helen and Milton A. Kimmelman Career Development Chair. O.A is the incumbent of The Miriam Berman Presidential Development Chair. O.A also acknowledges funding from the Henry Chanoch Krenter Institute for Biomedical Imaging and Genomics. Schemes in figures were prepared using BioRender.

## Authors contributions

A.G. and A.B.-S. designed the study and wrote the manuscript. A.G. carried out and analyzed all EV-related experiments, including EVs isolation, characterization, cell experiments, tumor inoculation, *in vivo* optical imaging and MRI experiments (in vitro and in vivo). H.A.-A. assisted with the establishment of the MRI experiments. J.Z. and G.S. designed the expression vectors of RBD and its variants, stablished the cell lines and purified the proteins. A.G. and J.Z. performed cell line characterization. P.A.K. and N.R.-R. guided and assisted with the establishment of EVs isolation, purification and characterization. M.I.M. and O.A. performed the cryo-TEM experiments. All authors reviewed the manuscript.

## Materials and Methods

### DNA manipulations

All DNA fragments were PCR amplified using KAPA HiFi HotStart ReadyMix (Roche, Switzerland) and purified by NucleoSpin® Gel and PCR Clean-up Kit (Nachery-Nagel, Germany). Restriction-free cloning PCR reactions (50 ul, KAPA HiFi HotStart ReadyMix) were composed of 100 – 200 ng of cloned DNA fragment and 20 ng of destination plasmid. The assembly PCR reactions consisted of 30 cycles of 1 min annealing, at 60 °C, and six mins of polymerization and 20 s at 98°C for denaturation. The template DNA was removed by using *DpnI* type IIM restriction enzyme (NEB, USA) at 37°C (1 – 2 h) and 0.9 μl from the reaction mix were electroporated to *E.coli* Cloni® 10G cells (Lucigen, USA). Colony PCR and sequencing were used for analysis and verification.

### pAGDisplay vector construction

pDisplay Mammalian Expression vector was purchased from Invitrogen (V66020). The pAGDisplay vector backbone was assembled by combining a pET28b fragment bearing KanR and origin of replication, a pLVX vector fragment bearing WPRE, PuroR, and IRES sequences, and a pDisplay CMV promoter with a PDGFRβ expression cassette by a restriction-free three-component assembly ^58^. In the subsequent restriction-free cloning step, the PuroR was fused with eUnaG2 fluorescent protein at the C-terminus ^59,60^. The full-length pAGDisplay was sequenced to verify its correct assembly.

### pAGDisplay modification with RFnano near-infrared fluorescence protein

To increase our spatial resolution, we introduced, by restriction-free cloning, a modified near-infrared miRFP670nano ^61^, an RFnano protein, as a cytoplasmic domain at the C-terminus of PDGFRβ. The preparation and design of RFnano is the subject of a publication currently in preparation. Briefly, the PROSS and Rosetta-based stabilization design was combined with S.cerevisiae EBY100 expression and sorting to obtain a brighter signal and the same spectral parameters. In total, 23 mutations were introduced into the protein.

### Protein production, purifications, and labeling procedures

The designed ALFA-tag binding nanobody (DnbALFA) and its mNeonGreen fusion were expressed by using expression plasmid pET28bdSUMO ^62^ and E.coli BL21(DE3) cells as described previously ^60^. Briefly, 200 ml of 2YT medium (16 g tryptone, 10 g yeast extract and 5 g NaCl, pH 7) was inoculated (1%), grown to the OD600 = 0.6 (37°C), and the expression was initiated by the addition of IPTG to a final concentration of 0.5 mM. The expression continued for the next 16 h at 20 °C. Expressed cells were washed (50 mM Tris-HCl, 200 mM NaCl buffer, pH 8), disintegrated by sonication, immobilized on Ni-NTA agarose (PureCube Ni-NTA Agarose, Cube Biotech, Germany), and cleaved on a column by using bdSUMO protease ^63^. The eluted fraction was further purified by size exclusion chromatography on HiLoad 26/600 Superdex 75 using a GE AKTA Purifier FPLC System. Soluble his-tagged peptidase domain of ACE2 protein (Q18 – S740), inserted in pHLsec plasmid, was expressed in an Expi293F cell system with an ExpiFectamine 293 Transfection Kit (ThermoFisher, USA) according to the manufacturer’s protocol and purified as described previously ^45^.The ACE2 or DnbALFA proteins were labeled by CF®640R succinimidyl ester dye (Biotium, USA, cat. 92108) in 0.1 M bicarbonate buffer using a 1:4 protein to dye ratio. The reaction was continued for 1 h at room temperature, and subsequently, the free dye was removed by dialysis (GeBAflex-Midi Dialysis Tubes, 8kDa MWCO, Geba, Israel) against PBS buffer (16 h, 4°C).

### Cell culture and transfection

HEK293, HEK293T, ACE2-expressing HEK293T cells, and stable pAGDisplay-RBD cell lines were cultured at 37°C in a 5% CO2 atmosphere in DMEM medium (4.5 g/L glucose, L-glutamine; Gibco, USA) supplemented with 10% fetal bovine serum (FBS) and 1% glutamine (4 mM). Cells were passaged 2-3 times per week using trypsin EDTA solution A (Biological Industries, USA) for cell detachment. The stably expressing ACE2 HEK293T cell line was kindly obtained from the lab of Dr. Ron Diskin (Weizmann Institute of Science) and kept under puromycin antibiotics (0.5 μg/ml, Invitrogen, USA).

### HEK-pAGDisplay-RBD stable cell line generation

The pAGDisplay-based plasmids expressing RBD (1 ug of DNA) were transfected in a 60 mm culture dish with 80% confluent HEK293 cells by a JetPrime transfection reagent (Polyplus, France) according to the manufacturer’s protocol. After transfection (24 h), cells were transferred to a 150 mm culture dish. Subsequently, 48 h after transfection, the media was replaced by fresh DMEM medium supplemented with 10% FBS and 1 μg/ml puromycin (Invitrogen, USA). Puromycin-resistant cells were selected for one week with regular replacement of cell media. Stably transfected cells associated with the top 1% green fluorescence signals (Puro-eUnaG2), were sorted out from the population using the S3e Cell Sorter device (Bio-Rad, USA) and further sub-cultured to single colonies.

### Characterization of parental cells

The presence of the RBD on the cell surface was measured by fluorescence-activated cell sorting (FACS). The cells were cultured until full confluence, then they were detached from the plates by PBS and added to the Eppendorf tubes. After spinning down (5 min at 500g), the cell pellet was resuspended in a labeling solution containing fluorescently labeled, purified ACE2 protein in PBS supplemented with 2% FBS. The cells were labeled for 1h on ice and then washed two times with 1 mL of PBS (spun down each time for 5 min at 500 g). After the last wash, the cells were resuspended in PBS with 2% FBS solution, filtered by 0.45 μm filters, and added to the FACS tubes. The fluorescence was measured by the LRSII FACS machine (BD Biosciences, USA) and analyzed by the FlowJo software.

### EV isolation

EV-depleted medium was prepared by two rounds of ultracentrifugation (100,000 g, 16 h) of DMEM with 20% FBS, diluted to 10% FBS and supplemented with glutamine. For EVs isolation, cells were cultured in EV-free medium for 48-72 h, then the medium was collected and processed by differential centrifugation (400 g, 10 min; 2000 g, 10 min; 10,000 g, 30 min, all at 4°C). The final supernatant was collected and EVs were pelleted at 100,000 g at 4°C for four hours in an Optima ultracentrifuge using the Beckman Ti45 rotor (Beckman Coulter, USA). The EVs pellet was washed with PBS and resuspended in 0.22 μm filtered PBS. EVs were isolated from the RBD-transfected cells (EV^RBD^) and control cells (EV^noRBD^).

### EV NTA analysis

The size and concentration of EVs diluted in PBS (1:1000 or 1:5000) was measured by nanoparticle tracking analysis using the NanoSight system (Malvern Instruments, UK) with a 405 nm laser by acquiring five, one-minute videos at the camera level 16. Threshold 5 was used for the analysis in all samples. Protein content was measured by the BSA (cells) or microBSA (EVs) protein assay according to the manufacturer’s instructions (Sigma Aldrich, USA).

### Western blot

For the Western blot analysis, EVs or parental cells were lysed by RIPA buffer (1X for cells, 1:1 in PBS for EVs) supplemented with a proteinase inhibitor cocktail. Proteins were separated on ExpressPlus™ PAGE Ready gels 4 - 20% (A2S, M42015) and transferred on the cellulose membrane (300 mA, 90 min) using a BioRad blotting device (BioRad, USA). After overnight blocking in 5% milk in TBST, specific antibodies were applied to the membranes for 1 h to detect markers: CD81 (B-II, Santa Cruz, USA); c-myc (9E10, Santa Cruz, USA) (all 1:500); and beta-actin (C4, Santa Cruz, USA) (1:1000). HRP-conjugated anti-mouse secondary HRP goat anti-mouse IgG antibody (#4053, Biolegend, USA) (1:5000 in TBST) was applied for 1 h before imaging using enhanced chemiluminescence substrate EZ-ECL Kit (Biological industries, catalog no. 20-500-120). An ALFA tag was detected by a homemade fluorescent anti-ALFA tag nanobody and visualized by a fluorescent reader at 642/662 nm wavelength.

### Binding assays and affinity curve determination using cell-display

The HEK-pAGDisplay-RBD stable cells grown to 80% confluency were gently detached by Accutase (1/2 solution in PBS, 3 min, Sigma Aldrich cat. A6964) and washed by PBS with 2 g/L BSA (PBSB). Aliquots of detached cells (approximately 2×10^6^) were incubated with a series of CF®640R succinimidyl ester-labeled (Biotium, USA, cat. 92108) ACE2 extracellular domain (AA Q18 – S740) solutions (0.2 – 83 nM) in 1 ml of PBSB for two hours on ice. After the incubation, cells in aliquots were separated by centrifugation (500 g, 5 min), washed, and resuspended in ice-cold PBSB, passed through a cell strainer nylon membrane (100 μM, SPL Life Sciences, Korea), and analyzed. The expression (eUnaG2-Puro signal, FL-1, excitation 498 nm, emission 527 nm) and binding (CF640R-ACE2, FL-4, excitation 642 nm, emission 662 nm) signals were recorded for 10k gated single-cell events by an S3e Cell Sorter (BioRad, USA). Mean FL-4 fluorescence signal values of RBD+ cells, subtracted by RFnano and the nonspecific signal of the RBD-population, were used to determine the KD of binding constants using a non-cooperative Hill equation and a nonlinear least-squares regression using Python 3.7 ^45^.

### Cryo-transmision electron microscopy (TEM)

#### Sample preparation

Lacey Carbon EM grids were glow-discharged (30 s, 25 mA) in a Pelco EasiGlow system. 3.5 μL of EVs resuspended in PBS -/- at ~10^11^ particles/mL concentration were applied onto the EM grid and the grid was incubated in the humidity chamber of Vitrobot Mark IV instrument (Thermo Fisher Scientific, USA) for 5 min at 100% humidity and at room temperature, followed by blotting (4.0 s and −10 Blot Force), and plunge-freezing into precooled liquid ethane. Samples were imaged using a Talos Arctica G3 TEM/STEM microscope (Thermo Fisher Scientific, USA), equipped with a OneView camera (Gatan, USA) at an accelerating voltage of 200 kV using SerialEM ^64^. Images were recorded at ×73000 magnification (calibrated pixel size 0.411 nm) with −3.5 μm defocus.

### *In vitro* binding of EVs^RBD^ to the ACE2 receptor

#### Flow cytometry

To confirm binding and uptake of EV^RBD^ in ACE2-expressing cells, isolated EVs^RBD^ or EV^noRBD^ were fluorescently labeled with 1,1’-Dioctadecyl-3,3,3’,3’-Tetramethylindotricarbocyanine Iodide (DiR) (ThermoFischer Scientific, D12731) at a concentration of 15 μM by incubation for 1h at room temperature. EVs were then washed three times with the VivaSpin centrifugation filters—five min at 14 000 g, and two min at 1000 g in a reverse position. HEK293T or ACE-expressing HEK293T cells were incubated with labeled EVs^RBD^ or EV^noRBD^ in EV-depleted DMEM for one or three h. Then, the cells were detached from the plate by PBS, washed twice with PBS in the Eppendorf tubes (spun down five min at 500 g), filtered by 0.45 μm filters, and added to the FACS tubes in FACS solution (PBS with 2% FBS). The fluorescence signal (APC-Cy7 filter) of cells was measured using a LSRII cell analyzer flow cytometer (BD Biosciences) and analyzed by the FlowJo software.

#### Fluorescence microscopy

For visualization by fluorescence microscopy, cells were seeded on 14 mm-diameter coverslips in a 24-well plate. The wells were coated with fibronectin by incubation for 45 min. Then, fibronectin was removed and the wells were washed with PBS before seeding the cells. For uptake of EVs in cells, the DiD-labeled EVs were dissolved in exosome-free medium and incubated with cells for three hours. Then, the medium was removed, and cells were washed two times with PBS. For staining of cell nuclei, DAPI was dissolved in 2.5% formaldehyde and the cells were incubated in the solution for 20 min. After two washes, the coverslips were carefully transferred and mounted on glass slides and the fluorescence was visualized using a wide-field microscope (Leica DMI8).

### Detection of ACE2 in cells

For confirmation of the presence of ACE2 protein in the ACE2-expressing cells, ACE2-expressing HEK293T and HEK293T control cells were lysed and the activity of the ACE protein was assessed by the SensoLyte 390 ACE2 Activity Assay Kit (AnaSpec, USA) according to the manufacturer’s instructions. The fluorescence (excitation 330 nm, emission 390 nm) of the final product was read within 5 - 35 minutes using a plate reader. The expression of ACE2 protein in cells was also confirmed by Western blot analysis, as described above, using a rabbit monoclonal ACE2 antibody (ab239924; 1:1000; Abcam, USA). For FACS analysis, the cells were seeded in a 6-well plate. Then the cells were detached from wells by PBS and incubated with ACE2 primary antibody (ab239924; 1:1000; Abcam, USA) for 1 hour on ice, washed with PBS and then incubated with anti-rabbit fluorescently-conjugated secondary antibody (Alexa Fluor 647 nm) for 1 hour on ice. The cells were then washed with PBS, filtered with 0.45 μm filters and resuspended in PBS with 2% FBS. The fluorescence signal (APC filter) of cells was measured by a FACS machine (LRS-II) and analyzed by the FlowJo software.

### Toxicity evaluation

For the evaluation of the toxicity of the vesicles, HEK293T cells were seeded in a 96-well plate a day before the addition of EVs^RBD^ or EVs^noRBD^. The cells were incubated with vesicles of different concentrations (1.8×10^8^ - 5×10^9^ EVs/well) for three h or 24 h, then the CellTiter-Blue solution was added to the cells according to the protocol (20 μL of CellTiter solution + 80 μL of culture medium). After incubation, the fluorescence of each well was read using a plate reader at excitation 560 nm and emission 590 nm (quadruplicates for each EVs concentration). The control sample contained no EVs.

### Production of magnetically labeled EVs

HEK293 cells stably expressing RBD or without RBD (control) were seeded into 15 cm culture plates at 50% confluency. The next day, SPIONs (Molday ION Dye Free; 2 mg Fe/mL; BioPal, USA, CL-50Q02-6A-0) were added to the culture medium at a final concentration of 40 μg/mL. Then, 24 hours after, medium was discarded, the cells were washed three times with 10 mL of PBS and EV-depleted medium was added to the cells followed by another 24-hour incubation. Afterward, EVs were isolated from the medium according to the standard protocol and resuspended in PBS.

### Animal model

Immunodeficient Hsd:Athymic Nude-Foxn1nu mice (Envigo) were used in the experiments. All animal studies were approved in accordance with the Weizmann Institute’s Animal Care and Use Committee (IACUC) guidelines and regulations (approval number 00580120-3). All animals were kept in a daily controlled room at the Weizmann Institute of Sciences animal facility with a surrounding relative humidity level of 50 ± 10% and a temperature of 22 ± 1 °C, with a 12/12 cycle of dark and light phases. Subcutaneous xenograft tumors were induced by injection of HEK293T (left side) and ACE2-expressing HEK293T cells (right side) under the skin above the mouse flank. Mice were kept under isoflurane anesthesia during the whole procedure. Each mouse received 10 mil of each cell type in 150 μL of PBS. The tumors were allowed to grow for two to three weeks until they reached a sufficient size for imaging (diameter around 0.5 – 1 cm). If the tumors reached more than 1 cm or the tumor size was not equal, the animals were removed from the experimental groups.

### Magnetic Resonance Imaging

MRI experiments were conducted on a horizontal 15.2 T horizontal scanner (*in vitro* and *in vivo*) ausing a ^1^H volume coil rf with a 23 mm diameter (Bruker BioSpin, Germany).

#### Phantom measurements

Different concentrations of isolated EVs were added to glass tubes and imaged using a RARE (Rapid Acquisition with Relaxation Enhancement) sequence with the following parameters: time of repetition (TR) 2800 ms; echo time (TE) 42 ms; and resolution 0.23×0.23×1 mm^3^. MR images were analyzed by the ImageJ software or by a custom-made script written in MATLAB (MathWorks, USA).

#### In vivo MRI of mice

Prior to MR imaging, a cannula was inserted into a mouse tail. Mice with induced tumors were then measured on a 15.2T MRI scanner before and after injection of SPIONs-labeled EVs^RBD^ at a dose of 3×10^11^ EVs in 100 μL of PBS administered through a long tube without changing the position of the animal. The mice were measured under general isoflurane inhalation anesthesia (5% induction, 1% maintenance) up to four hours after EVs injection using a gradient echo Fast Low Angle Shot (FLASH) with the following parameters: TR = 300 ms; TE = 2 ms; resolution of 0.23×0.23×0.7 mm^3^. After tuning and matching to the^1^H frequency, shimming of the magnetic field, and B_0_ correction, the tumor area was measured with the following parameters: both axial and coronal images were acquired; T2 and T2* maps were reconstructed in the Paravision software; and were followed by analysis in the ImageJ software. Regions of interest (ROI) were outlined around each tumor, muscle, and a noise area outside of the animals. The contrast-to-noise ratios (CNR) were calculated as follows:

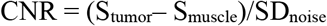

CNR difference was calculated as a percentage of CNR before and after EV injection.

### Fluorescence imaging

Prior to imaging, mice with induced subcutaneous tumors were retro-orbitally injected with DiR-labeled EVs^RBD^ or EVs^noRBD^ (dose of 3×10^11^ EVs in 100 μL of PBS). Six hours after injection, the mice were intracardially perfused with 2.5% formaldehyde solution under general anesthesia induced by ketamine (80 mg/kg) and medetomidine (0.6 mg/kg). Organs (liver, kidneys, spleen, intestine, heart, lungs, brain) and tumors were excised, fixed with 2.5% formaldehyde solution overnight, and then kept in PBS. A day after, fixed organs were measured with IVIS Lumina XR optical images (Perkin Elmer, USA) with the FOV and exposure time of 1 s (liver), 10 s (kidney, lungs, spleen, heart), 30 s (brain), or 60 s (tumors). For intravital microscopy, an Olympus microscope MVX was used with exposure times of 700 ms (Cy7 filter for detection of the DiR signal).

### Histology

The fixed tumor tissues were kept in 1% formaldehyde solutions and then embedded in paraffin blocks according to a standard protocol. Tissue slices were cut on a microtome and stained by hematoxylin&eosin and Prussian blue. The slides were imaged on a Leica DMI8 wide-field fluorescent microscope using a standard bright-field filter.

### Statistical analysis

All numerical data are presented as mean±standard deviation (s.d.). Statistical analysis was performed using GraphPad Prism 8.0 software (GraphPad Software Inc., USA). Comparison of two groups was analyzed by a two-tailed Student’s t-test. A p-value of 0.05 and below was considered significant: *p-value<0.05, ** p-value <0.01, ***, p-value <0.001 and **** p-value <0.0001.

