## Supplementary Information for "Genetically engineered MRI-trackable extracellular vesicles as SARS-CoV-2 mimetics for mapping ACE2 binding *in vivo*"

### Supplementary Figures

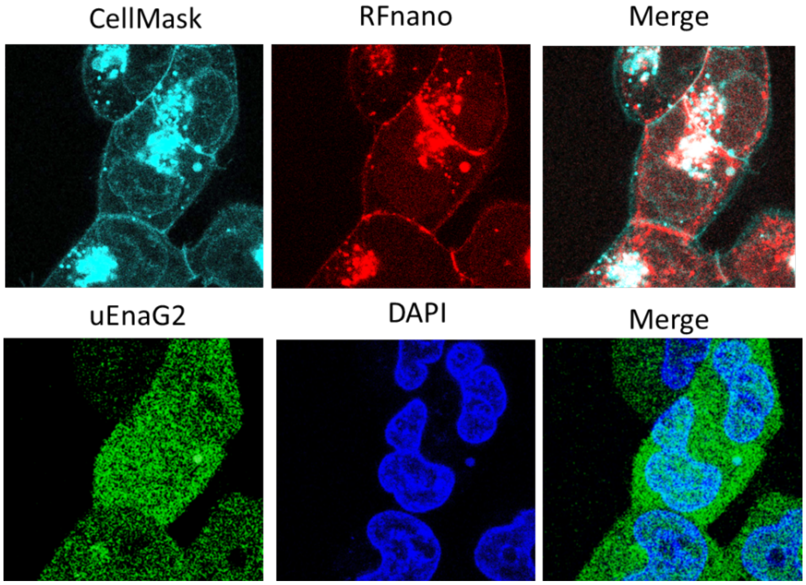


**Figure S1. Confocal microscopy of RBD cells.** A cell mask to label cell membranes was co-localized with miRFP670 fused with RBD, confirming membrane expression of RBD. The UnaG2 protein used for FACS sorting and establishment of a stable cell line was localized in cytoplasm.

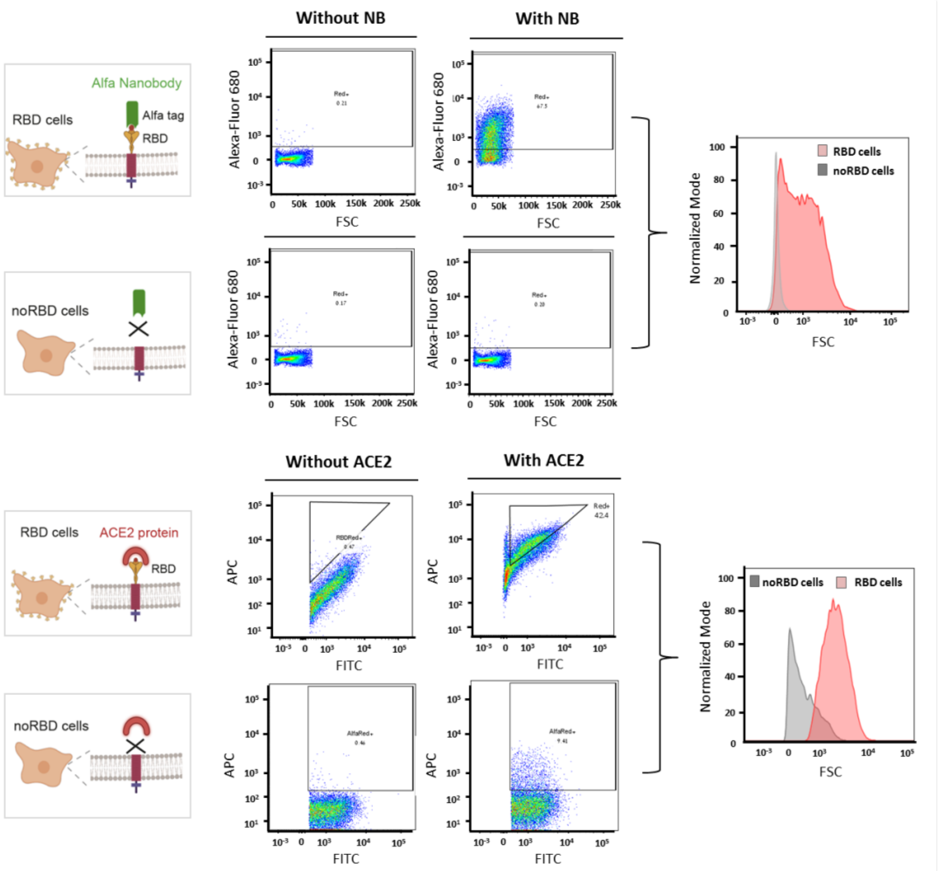


**Figure S2: Detection of ALFA tag (upper panel) and binding to ACE2 protein (bottom panel) in parental RBD/noRBD cells by flow cytometry.** Gating strategies for FACS.

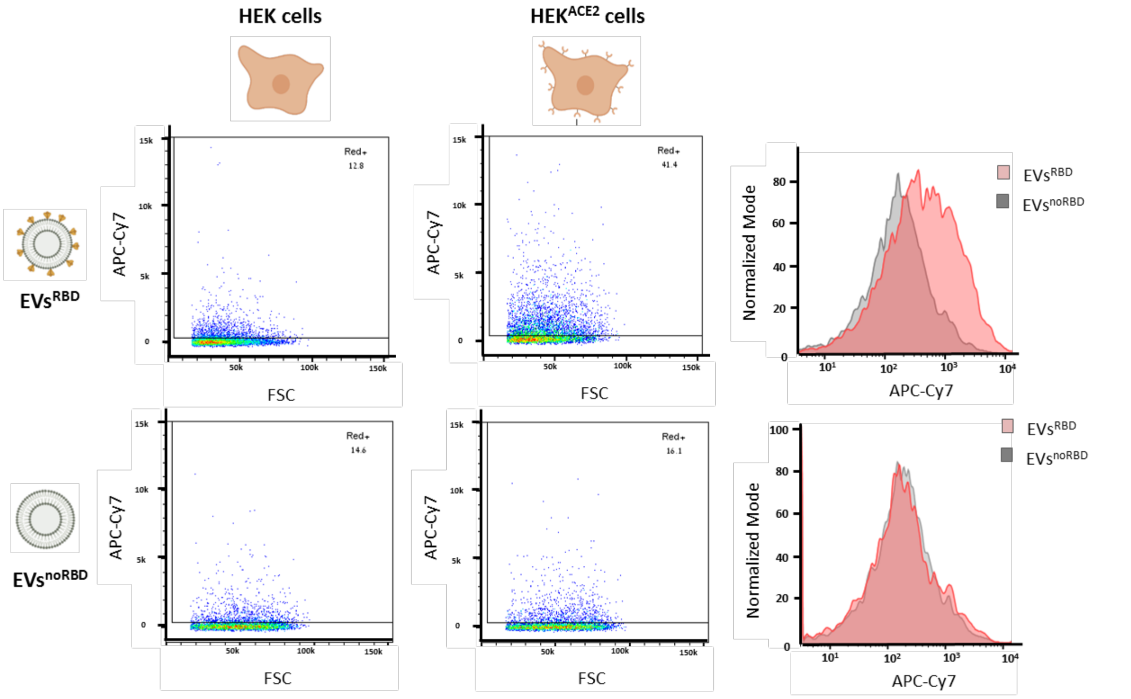


**Figure S3: Targeted uptake of EVs in cells.** Flow cytometry analysis of DiR-labeled EVs$^{RBD}$ and EVs$^{noRBD}$ accumulated in HEK (control) and ACE2-expressing cells—dot blots (left) and histograms (right).

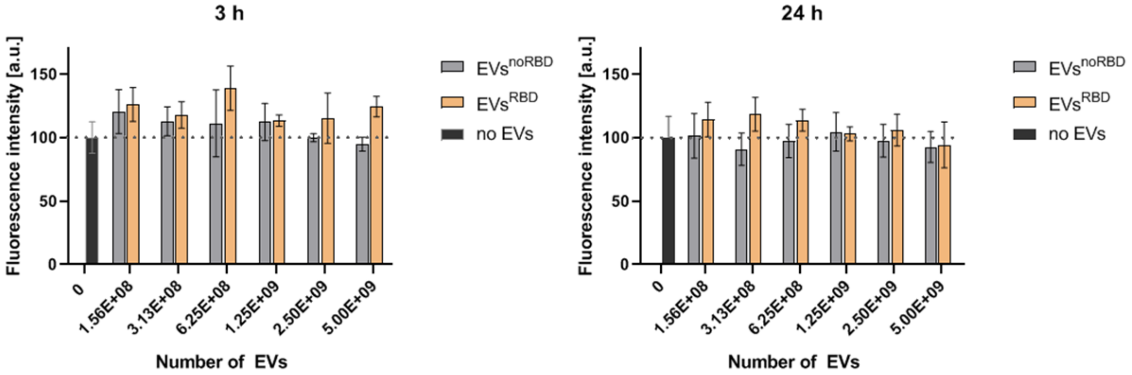


**Figure S4. Toxicity evaluation of cells with incubated EVs.** Cell-Titer-Blue assay of HEK293T cells incubated with EVs for 3 h (left) and 24 h (right).

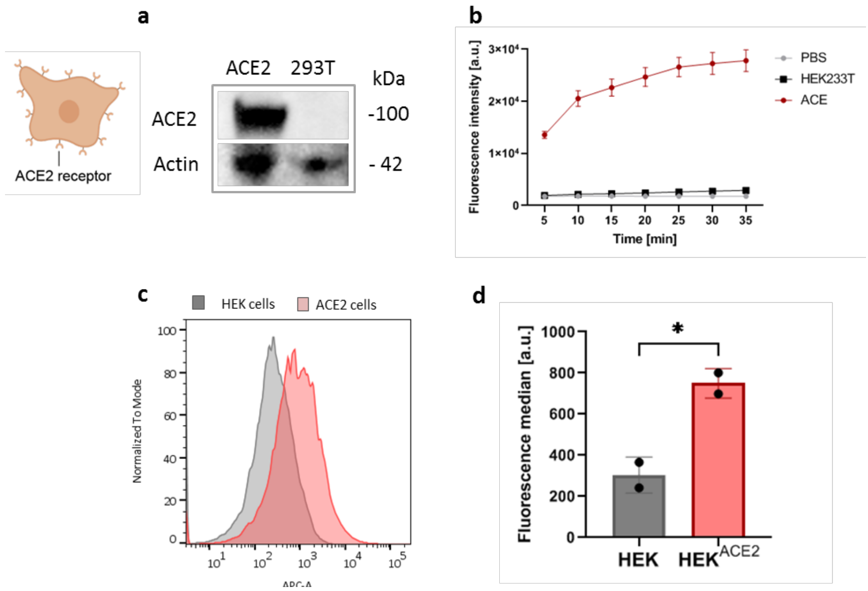


**Figure S5. Confirmation of ACE2 in ACE2-expressing stable cell line.** Western blot (a), enzymatic detection (b) and FACS analysis of HEK and HEK$^{ACE2}$ cells shown as (c) histogram and (d) quantification of fluorescence median.

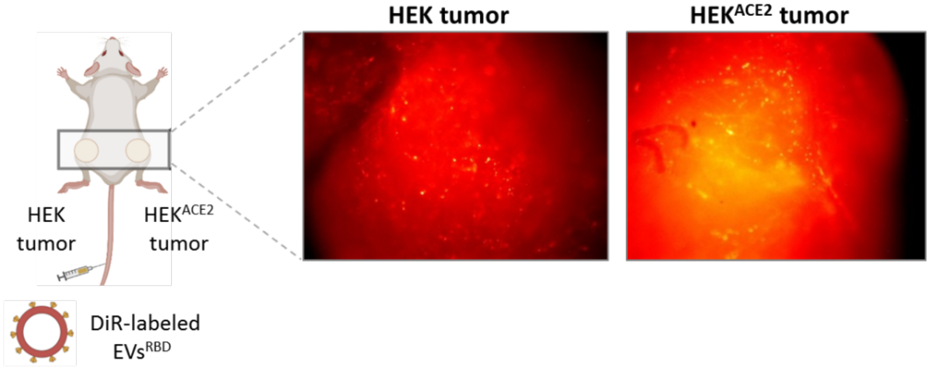


**Figure S6. Accumulation of EVs$^{RBD}$ in mice.** Intravital microscopy of tumor tissue showing individual EVs accumulated in the tumors.

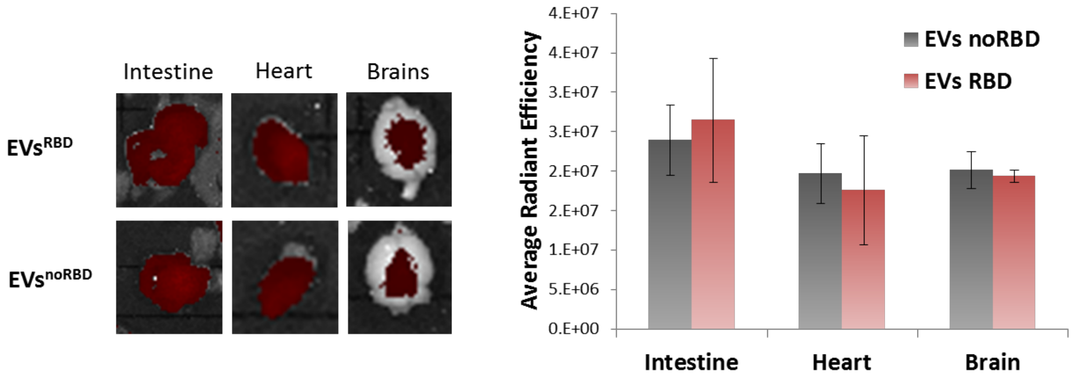


**Figure S7.** Biodistribution of EVs$^{noRBD}$ and EVs$^{RBD}$ in intestine, heart, and brain of mice.

**Supplementary sequences:**

**RBD-Wuhan**
TGCCCTTTTGGTGAAGTTTTTAACGCCACCAGGTTTGCCTCTGTCTATGCCTGGAACA
GGAAGAGGATTAGCAACTGTGTGGCTGACTACTCTGTGCTCTACAACTCTGCC
TCCTTCAGCACCTTCAAGTGTTATGGAGTGAGCCCAACCAAACTGAATGACCTGTGT
TTCACCAATGTCTATGCTGACTCCTTTGTGATTAGGGGAGATGAGGTGAGACAG
ATTGCCCCTGGACAAACAGGCAAGATTGCTGACTACAACTACAAACTGCCTGATGAC
TTCACAGGCTGTGTGATTGCCTGGAACAGCAACAACCTGGACAGCAAGGTGGGA
GGCAACTACAACTACCTCTACAGACTGTTCAGGAAGAGCAACCTGAAACCATTTGAG
AGGGACATCAGCACAGAGATTTACCAGGCTGGCAGCACACCATGTAATGGAGTG
GAGGGCTTCAACTGTTACTTTCCACTCCAATCCTATGGCTTCCAACCAACCAATGGAG
TGGGCTACCAACCATACAGGGTGGTGGTGCTGTCCTTTGAACTGCTCCATGCC
CCTGCCACAGTGTGTGGACCAAAG

**RBD-Delta**
TGCCCTTTTGGTGAAGTTTTTAACGCCACCAGGTTTGCCTCTGTCTATGCCTGGAACA
GGAAGAGGATTAGCAACTGTGTGGCTGACTACTCTGTGCTCTACAACTCTGCC
TCCTTCAGCACCTTCAAGTGTTATGGAGTGAGCCCAACCAAACTGAATGACCTGTGT
TTCACCAATGTCTATGCTGACTCCTTTGTGATTAGGGGAGATGAGGTGAGACAG
ATTGCCCCTGGACAAACAGGCAAGATTGCTGACTACAACTACAAACTGCCTGATGAC
TTCACAGGCTGTGTGATTGCCTGGAACAGCAACAACCTGGACAGCAAGGTGGGA
GGCAACTACAACTACAGATACAGACTGTTCAGGAAGAGCAACCTGAAACCATTTGA
GAGGGACATCAGCACAGAGATTTACCAGGCTGGCAGCAAACCATGTAATGGAGTG
GAGGGCTTCAACTGTTACTTTCCACTCCAATCCTATGGCTTCCAACCAACCAATGGAG
TGGGCTACCAACCATACAGGGTGGTGGTGCTGTCCTTTGAACTGCTCCATGCC
CCTGCCACAGTGTGTGGACCAAAG

**RBD-Omicron (version_3S)**
TGCCCTTTTGGTGAAGTTTTTAACGCCACCAGGTTTGCCTCTGTCTATGCCTGGAACA
GGAAGAGGATTAGCAACTGTGTGGCTGACTACTCTGTGCTCTACAACTCTGCC
TCCTTCAGCACCTTCAAGTGTTATGGAGTGAGCCCAACCAAACTGAATGACCTGTGT
TTCACCAATGTCTATGCTGACTCCTTTGTGATTAGGGGAGATGAGGTGAGACAG
ATTGCCCCTGGACAAACAGGCAATATTGCTGACTACAACTACAAACTGCCTGATGAC
TTCACAGGCTGTGTGATTGCCTGGAACAGCAACAAACTGGACAGCAAGGTGTCT
GGCAACTACAACTACCTCTACAGACTGTTCAGGAAGAGCAACCTGAAACCATTTGAG
AGGGACATCAGCACAGAGATTTACCAGGCTGGCAACAAACCATGTAATGGAGTG
GCCGGCTTCAACTGTTACTTTCCACTCAAGTCCTATTCTTTCCGTCCAACCTACGGAG
TGGGCCATCAACCATACAGGGTGGTGGTGCTGTCCTTTGAACTGCTCCATGCC
CCTGCCACAGTGTGTGGACCAAAG
